# Prey choice of the common vampire bat on introduced species in an Atlantic forest land-bridge island

**DOI:** 10.1101/2020.04.02.022236

**Authors:** Fernando Gonçalves, Marcelo Magioli, Ricardo S. Bovendorp, Katia Maria Paschoaletto Micchi de Barros Ferraz, Letícia Bulascoschi Cagnoni, Marcelo Zacharias Moreira, Mauro Galetti

## Abstract

The proliferation of native, alien, invasive and domestic species provide novel and abundant food resources for the common vampire bat (Desmodus rotundus) that could alter its prey preference. Based on the analysis of carbon and nitrogen stable isotopes, we report the prey choice of D. rotundus on introduced mammals in an tropical land-bridge island where the domestic animals were removed and 100 individuals of 15 mammal species were intentionally introduced. Our analysis shows that, D. rotundus on Anchieta Island were more likely to prey upon species from open habitats (mean value of −14.8‰), i.e., animals with high δ13C values characterized by the consumption of C4 resources. As expected for a top predator species, δ15N values for D. rotundus were higher (mean value of 8.2‰) and overlapped the niche of the capybaras (Hydrochoerus hydrochaeris) from the Anchieta Island, while it was distant from coatis, and also from those potential prey from the preserved area in the mainland, including the capybaras, indicating that among all potential mammalian prey species, they fed exclusively on capybaras, the highest mammalian biomass on island. Based on previous information on human occupation, the domestic animals present on Anchieta island might be the main prey of D. rotundus and responsible for maintaining a viable population. As the capybaras were introduced only 36 years ago, this suggests a rapid prey shift due to anthropogenic disturbances, which has allowed common vampire bats to successfully exploit them. Literature records also show that common vampire bats were not captured in preserved areas of the mainland which are near Anchieta Island indicating that the percentage of capture of D. rotundus is usually low in natural forested habitats where potential prey are scattered. As three individuals of introduced capybaras were confirmed died from bat rabies viruses (RABV) in 2020, we suggest periodic monitoring of bat rabies viruses in common vampire bat populations on Anchieta Island and areas nearby, in order to quantify the magnitude of the outbreak area and develop strategies for controlling, especially considering that the island and areas nearby is frequently visited by tourists. We highlighted that this prey choice is context-dependent, and possibly influenced by the removal of domestic animals, the explosive population growth of introduced capybaras combined with their predictable foraging behavior.

## Introduction

The common vampire bat *Desmodus rotundus* (Geoffroy, 1810), an obligate blood-feeding species and the primary reservoir of rabies, has experienced changing availability of both wild and domestic prey (Greenhall *et al.*, 1983; Galetti *et al.*, 2016; Gnocchi and Srbek-Araujo, 2017; Zortéa *et al.*, 2018) throughout its range from Mexico to northern Argentina. Usually, common vampire bats have low densities in old-growth forest (Bernard, 2001; Bobrowiec *et al.*, 2014; Gonçalves *et al.*, 2017) where potential prey are sparse, but their population increases in fragments surrounded by pastures (Delpietro *et al.*, 1992; Bobrowiec, 2012) due the increased availability of livestock species (Greenhall, 1988; Delpietro *et al.*, 1992; Bobrowiec, 2015). The higher livestock densities in the Neotropical region combined with introduction of native, alien and invasive species, has created a novel, abundant and reliable source of blood for common vampire bats, causing population growth and geographic range expansions (Delpietro *et al.*, 1992; Lee *et al.*, 2012; Bobrowiec *et al.*, 2015; Galetti *et al.*, 2016).

Detailed analyses of prey choice of common vampire bat in this anthropogenic scenario are fundamental to answer questions on trophic interactions, how predators and prey interact, and how prey availability affects predator density and distribution (Sheppard and Harwood, 2005). In the past two decades, studies have used stable isotope analysis and molecular typing of DNA in vampire bat faeces to demonstrate reliance on livestock when they are locally abundant (Voigt and Kelm, 2006; Bobrowiec *et al.*, 2015) and studies with camera traps based on video footage have revealed behavioral aspects of feeding on wild species (Castellanos and Banegas, 2015; Galetti *et al.*, 2016; Gnocchi and Srbek-Araujo, 2017; Zortéa *et al.*, 2018). However, prey choice in regions with introduced species rather than livestock have not been studied yet, but are of critical importance due to risks to public health and consequences for the transmission of infectious diseases by altering demographic processes, animal interactions and host immunity (Schneider *et al.*, 2009; Stoner-Duncan *et al.*, 2014; Streicker and Allgeier, 2016).

Here, we report, based on analysis of stable carbon and nitrogen isotopes, the prey choice of common vampire bats (*Desmodus rotundus*) on introduced mammals on a tropical island where 100 individuals of 15 mammal species were intentionally introduced 36 years ago. Our analysis shows that, between two suitable species classified as potential prey, they fed exclusively on capybaras (*Hydrochoerus hydrochaeris*), the highest mammalian biomass on the island. We highlight that this prey choice of common vampire bats are context-dependent, and possibly influenced by the removal of domestic animals, the explosive population growth of introduced capybaras combined with their predictable foraging behavior.

## Materials and Methods

### Study area

The study was carried out on Anchieta Island (23°27’S; 45°02’W), an 828-ha land-bridge island in Ubatuba, north coast of São Paulo State, Brazil (Fig. 1). The island is 500-meter away from the mainland and has a long history of human occupation and was called as Ilha dos porcos (Pigs Island) in allusion to the large number of pigs that existed on island (Guillaumon *et al.*, 1989). In the beginning of the last century, the island had a prison and, especially during the years when the prison was active (1904-1955), cattle, pigs, dogs, cats, and the domestic fowl were brought to the island (Galetti *et al.* 2009) in order to sustain its human community that reached over 1,000 residents belonging to 420 families (Guillaumon *et al.*, 1989). The prison and all infrastructures were expropriated and the island was transformed into a state park in 1977, and all the domestic animals were removed (Guillaumon *et al.*, 1989).

**Fig. 1:**
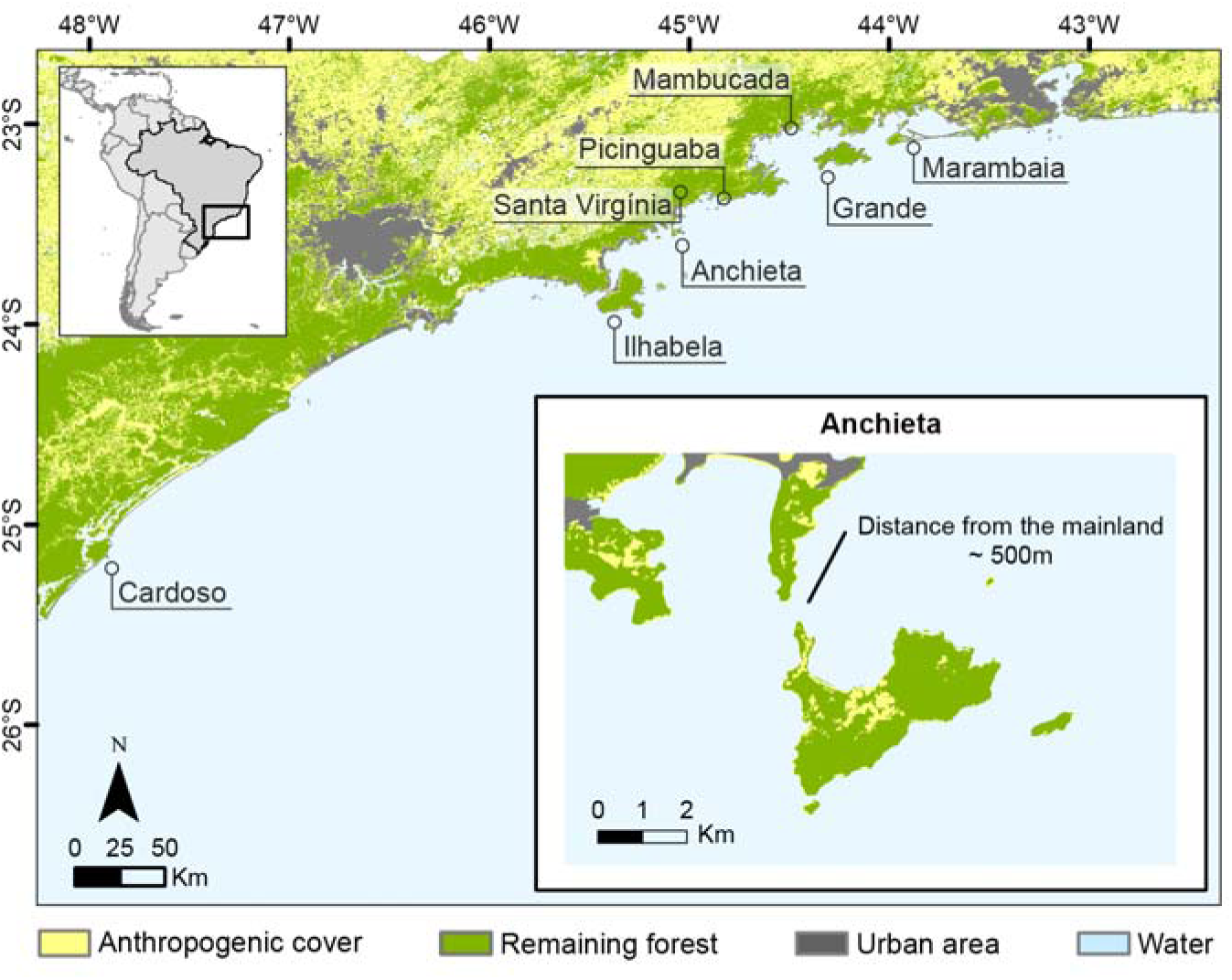
Location of Anchieta Island, state of São Paulo, southeastern Brazil and its proximity with another islands and mainland sites.

In March 1983, the São Paulo Zoo Foundation introduced 100 individuals of 14 mammal species on island (Guillaumon *et al.*, 1989). Of the 14 species introduced onto 1983, seven species have not been recorded since and have probably been extirpated (Bovendorp and Galetti, 2007; Supporting information S1). Little is known about the mammalian fauna on the island prior to human occupation. However, it is probable that it was similar to that on the continent given its proximity (500 m), with the exception of large predators (e.g., jaguars and pumas) and ungulates. Thirty-six years after introduction, the island now harbors the highest density of terrestrial mammals (486 ind./km^2^) in the entire Atlantic Forest (Bovendorp and Galetti, 2007). The vegetation on the island is composed of coastal Atlantic rainforest, with coastal plains where notable features include a stretch of *restinga* (a distinct type of coastal tropical and subtropical moist broadleaf forest), and large areas of disturbed vegetation dominated by ferns (*Gleichenia*).

In October 2017 and November 2018, we did two field expeditions in order to capture and sample the predator-prey systems. We selected three accessible forest sites, and three accessible open sites. Sample sites were selected without prior knowledge of the presence of common vampire bats or their potential prey. During each field expedition, each habitat was sampled for three consecutive days in order to equally divide efforts.

### Capture and sampling of common vampire bats

We mist-netted bats during three consecutive nights in two field expeditions. Netting was undertaken during the absence of moonlight, using 2.6 × 12 m ground-level mist nets opened for six hours after dusk. The number of nets per night ranged from three to six, but did not vary among sampling habitats (open area and forest). The capture effort (net area multiplied by the number of hours nets were open) in each habitat was 310 m^2^h. All captured bats were kept individually in cloth bags for 45–60 min, during which time we collected hair from the dorsal posterior region for stable isotope analysis. We then released the specimens at the site of capture.

### Capture and sampling of potential prey

According to literature records, common vampire bats just feed on medium-size and/or large mammals (> 1 kg), thus coatis (*Nasua nasua*) and capybaras (*H. hydrochaeris*) are the only suitable potential prey on the Island. In order to capture and collect hair from coatis, we used 30 live traps in each sampling site (forest and open), during three consecutive nights simultaneously to the mist nets, which resulted in an effort of 360 trap-nights. For capybaras, we collected hair samples stuck in barbed-wire fences in all trails on Anchieta Island. To complement our sampling effort, hair samples from coati and capybaras were collected opportunistically with tweezers. All captures, handling, and tagging techniques followed the guidelines of the Mammal Society (Sikes *et al.*, 2016).

To understand if the common vampire bat uses the mainland nearby Anchieta Island to feed on wild animals, we used isotopic values obtained by Magioli et al. (2019) on five large-bodied mammals [white-lipped peccary (*Tayassu pecari*), collared peccary (*Pecari tajacu*), deer (*Mazama* sp.), lowland tapir (*Tapirus terrestris*) and capybara] from a protected area of the mainland (Núcleo Santa Virgínia, an administrative division of the Serra do Mar State Park). This area is inserted in the largest continuous remnant of the Atlantic Forest, and distance ∼19 km (Fig. 1) in a straight line from Anchieta Island.

### Common vampire bats feeding on potential prey

We observed common vampire bats feeding by *ad libitum* sampling (Martin and Bateson, 2007) during 17 nights (47 hours of observation), in October 2017 and November 2018 in the same 6 selected sampling sites. Observations occurred between specific shifts of two to five hours per night, between 6pm and 5am. Observers, equipped with red flashlights, were situated on high ground in order to see the entire area. When capybara were detected, the observers approached them slowly, and observed if common vampire bats were feeding. If feeding was detected, we recorded it (WebVideos S1 and S2).

### Data analysis

We also compiled bat capture data from literature, including capture effort, from only two previous studies on the island (Aires, 1998; Colas-Rosas, 2009), in order to complement our species list and to estimate the percentage of capture. The percentage of capture of common vampire bats on Anchieta Island was calculated by dividing the number of total common vampire bat mist-netted by the effort (m^2^h) multiplied by 100. We used data from Bovendorp and Galetti (2007) to estimate the potential prey density on the island, and to estimate prey biomass, we used body mass data from Gonçalves et al. (2018).

### Stable isotopes analysis

To analyze the stable carbon and nitrogen isotopes, we cleaned the hair samples with water and 70% alcohol to remove any residue and dried them with absorbent paper. We then cut up the samples and stored them in thin capsules. Later, we used a CHN-1110 Elemental Analyzer (Carlo Erba, Milan, Italy) to combust the material, and separated the resultant gases in a chromatographic column. Lastly, we inserted the gases in a coupled continuous flow isotope ratio mass spectrometer (Delta Plus, Thermo Scientific, Bremen, Germany) to obtain the isotopic composition of the samples. The isotopic values of carbon and nitrogen were expressed in delta notation (δ^13^C, δ^15^N) in parts per mil (‰) relative to the V-PDB (Vienna-Pee Dee Belemnite) and atmospheric N_2_ standards, respectively. Delta values were calculated based on the standards using the following equation δX = [(R_sample_**/**R_standard_) ◺ 1] multiplied by 1000, where X represents the stable carbon or nitrogen isotopes (^13^C or ^15^N), and R the isotopes ratio (^13^C/^12^C or ^15^N/^14^N).

We performed the replication of the same individual material for only 10% of the samples, but the precision of the analytic method for 22 replicas of an internal standard for all batches, was estimated as 0.09‰ for carbon and nitrogen. The samples were anchored to international scales by the use of international reference materials: NBS-19 and NBS-22 for carbon, and IAEA-N1 and IAEA-N2 for nitrogen.

### Resource use

To obtain information on resource use of common vampire bats, we adapted the analytical approach used by Magioli et al. (2014, 2019). The analysis consists of using a simple mixed model that interpolates the stable carbon isotopic values of samples, accounting for specific fractionation factors, with the mean values of the different vegetation types (C_3_ and C_4_ plant photosynthetic cycles), while also considering the minimum and maximum values obtained for all animal samples analyzed. To estimate fractionation factors (Δ^13^C and Δ^15^N), we used the ‘SIDER’ package (Healy et al. 2018), available in R 3.5.3 (R Development Core Team 2019), that estimates species-specific fractionation factors from phylogenetic regression models, accounting for a database of fractionation values available for several species. We generated the Δ^13^C value for the common vampire bats (2.1 ± 1.9‰) using the script available in (Healy *et al.*, 2018).

To determine the origin of food items consumed by the mammals, i.e., C_3_ or C_4_ plants, we calculated the carbon content in each sample (δ^13^C values corrected by Δ^13^C values) using the following equation:

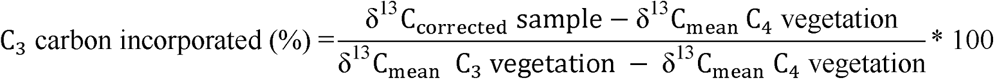

We used as base for our model, the mean δ^13^C value of −32‰ to indicate C_3_ plants, and −12‰ V-PDB to C_4_ plants. These values were obtained from the extreme δ^13^C_corrected_ values of all animal samples analyzed (predator and prey). After calculating the proportion of C_3_/C_4_ carbon, we classified samples in three groups: (1) C_3_ group – species that preferentially consumed C_3_ items (> 70% of C_3_ carbon; δ^13^C = −32 to −26‰); (2) Mixed group – species that used both C_3_ and C_4_ food items (from 30 to 70% of C_3_ carbon; δ^13^C = −25.9 to −18.1‰); C_4_ group – species that mainly consumed C_4_ items (< 30% of C_3_ carbon; δ^13^C = −18 to −12‰). We also corrected the δ^15^N values using the fractionation factor (Δ^15^N = 3.4 ± 1.5‰) generated by the ‘SIDER’ package.

### Isotopic niches

To assess the overlap of resource use by the common vampire bat and its potential prey, we analyzed the size of the isotopic niches using the ‘SIBER’ package (Jackson *et al.*, 2011), available in R 3.5.3. This package calculates the standard ellipses area (SEA) using δ^13^C_corrected_ and δ^15^N_corrected_ values, which contain 95% of the data, independent of sample size, allowing comparison of the isotopic niche width between species. To control sample size, we used the SEA corrected (SEAc).

## Results

### Capture of common vampire bats and potential prey

We recorded 187 individuals (16 of common vampire bats) belonging 13 bat species on Anchieta Island (Supporting information S2) and collected fur of 17 individuals of capybaras and 10 coatis (Supporting information S3 and S4). The percentage of capture for common vampire bats was 0.12% (16 individuals/12607 m^2^h * 100) (Supporting information S2), while the mean density of both potential prey was estimated as 60.9 individuals/km^2^ (coati = 25.06 and capybara = 35.30) (Table 1). Capybaras showed the highest mean biomass (1,112 kg/km^2^) on the island (Table 1). Due to the predictable foraging behavior of capybaras in open areas, only the bat-capybara system was detected by observers. The common vampire bat fed on capybaras in 17 observations during 47 hours of sampling effort (Supporting information S5, WebVideos S1 and S2).

### Prey choice of the common vampire bat

Common vampire bats on Anchieta Island were more likely to prey upon species from open habitats (mean value of −14.8‰), i.e., animals with high δ^13^C values characterized by the consumption of C_4_ resources. The δ^15^N values for common vampire bats were higher than expected and most likely similar with apex predator species (mean value of 8.2‰) (Fig. 2, Supporting information S3). One of the potential prey – the coati – largely depended on resources from the forest remnants (C_3_ resources) (Fig. 2). Capybaras presented a large isotopic niche, using resources from both open areas and forest remnants, but feeding mainly on C_4_ plants (Fig. 2). The isotopic niche of common vampire bats overlapped the niche of the capybaras, while it was distant from coatis (Fig. 2), and distinct from mean values of potential prey in the mainland, including the capybaras there (Fig. 3).

**Fig. 2:**
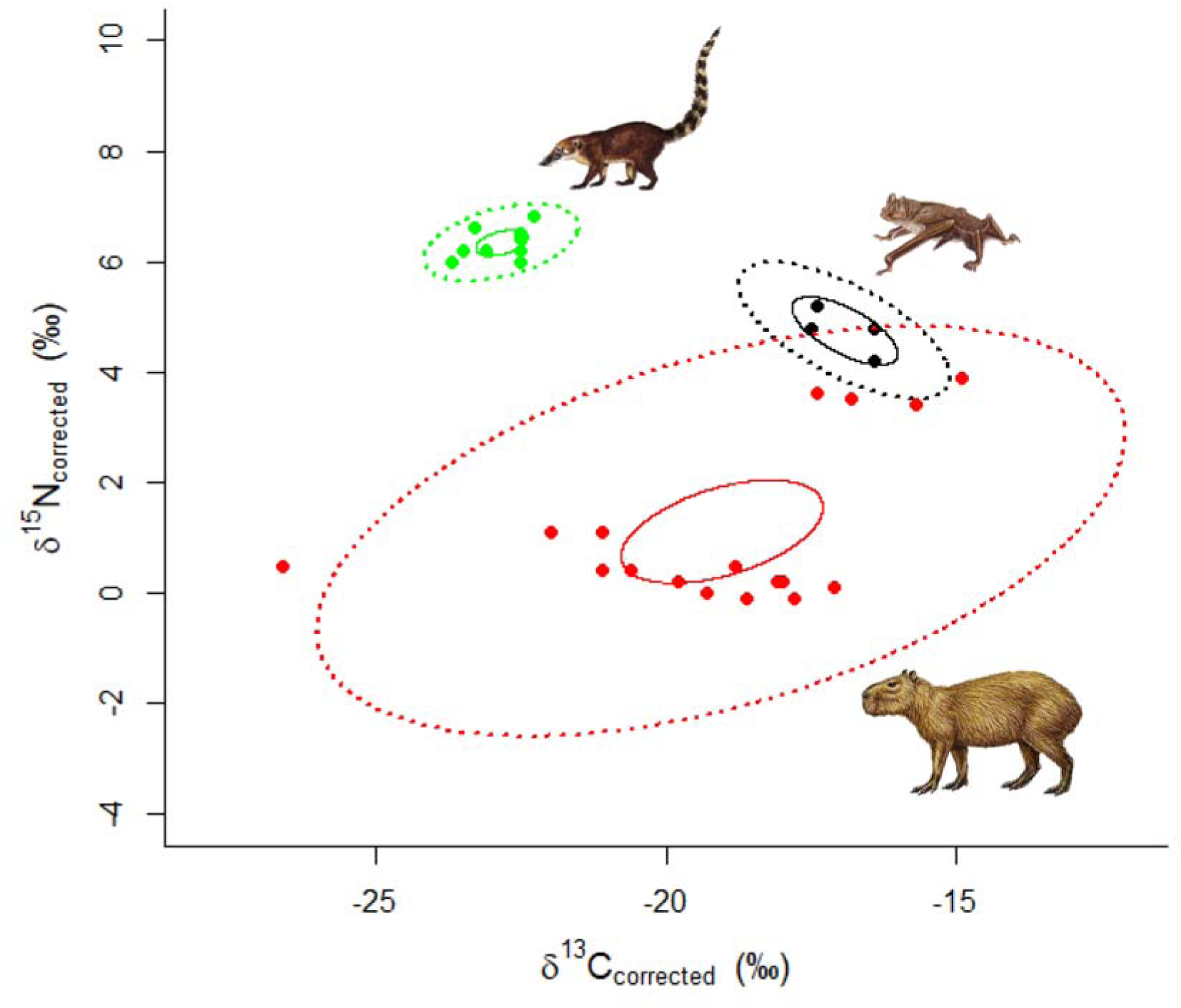
Isotopic niches (standard ellipses area corrected - SEAc) and individual values (δ13C, δ15N) of common vampire bats (Desmodus rotundus) (black) and its potential prey on Anchieta Island, state of São Paulo, southeastern Brazil. Isotopic values for D. rotundus were corrected using species-specific fractionation factors (Δ13C = 2.1‰; Δ15N = 3.4‰). Dashed lines = estimated standard ellipses using 95% of the data; solid lines = confidence intervals (95%) around the bivariate means; capybara (Hydrochoerus hydrochaeris) = red; Coati (Nasua nasua) = green.

**Fig. 3:**
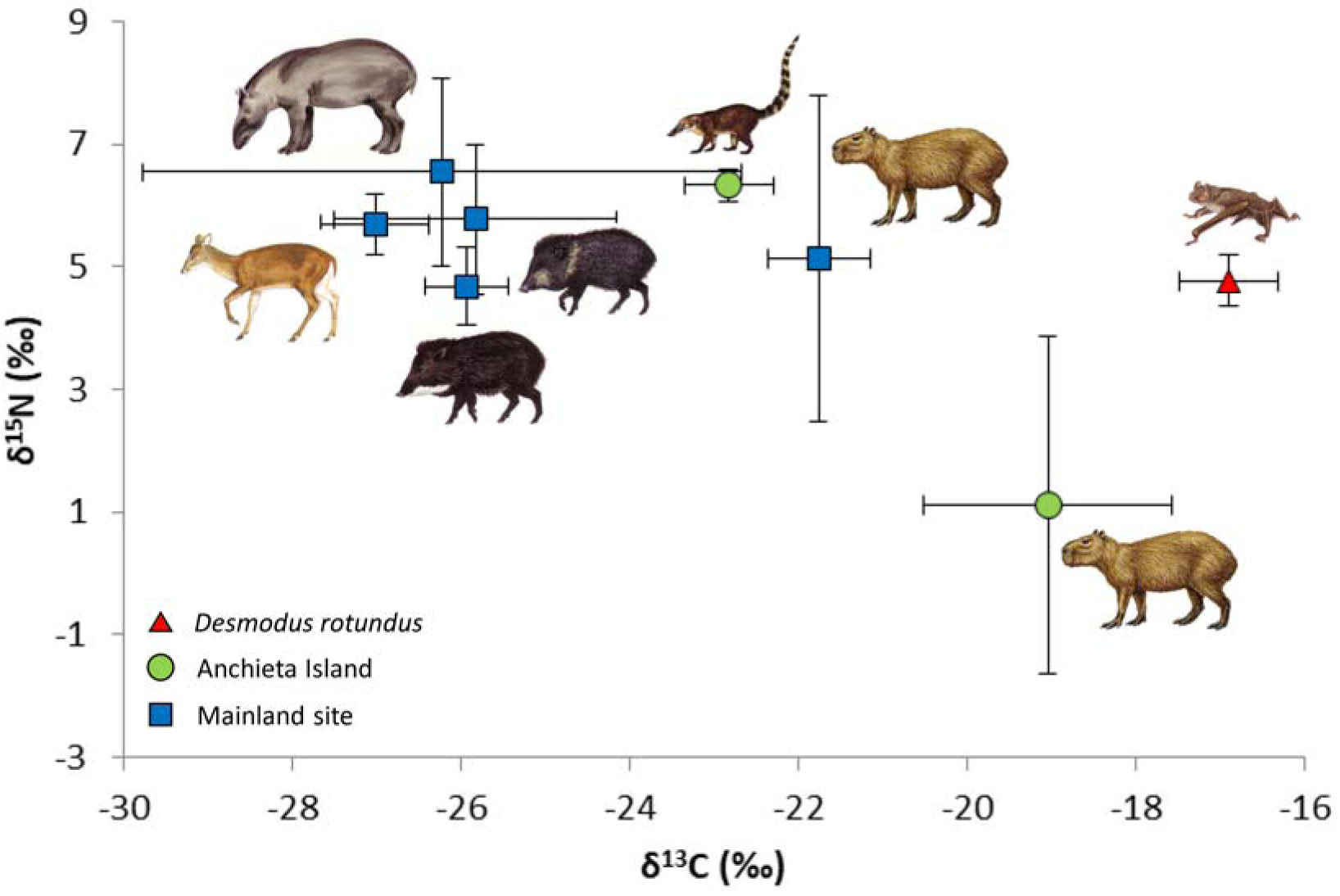
Mean and standard deviation of δ13C and δ15N values of common vampire bats (Desmodus rotundus; red triangle) and its potential prey on Anchieta Island (green circles) and Núcleo Santa Virgínia of the Serra do Mar State Park (mainland site; blue squares), state of São Paulo, southeastern Brazil. Isotopic values for D. rotundus were corrected using species-specific fractionation factors (Δ13C = 2.1‰; Δ15N = 3.4‰). Anchieta Island potential prey: Capybara (Hydrochoerus hydrochaeris) and coati (Nasua nasua). Mainland site potential prey: White-lipped peccary (Tayassu pecari), collared peccary (Pecari tajacu), deer (Mazama sp.), lowland tapir (Tapirus terrestris) and capybara.

## Discussion

The isotopic niche of common vampire bats overlapped the capybaras’ niche from the Anchieta Island (Fig. 2), and was distinct from the mean values of potential prey in the preserved area of the mainland (Fig. 3), indicating that capybaras from the Anchieta island are their main food source. Even with concentrated sampling effort, our results support previous studies that showed a choice of common vampire bats to feed on locally abundant and reliable prey (Voigt and Kelm, 2006; Bobrowiec *et al.*, 2015; Streicker and Allgeier, 2016, Zórtea *et al.*, 2018). Previous long-term isotopic studies, which analyzed tissues with different isotopic turnover rates (e.g. blood, skin, hair from the different individuals and assemblages) showed that the common vampire bat dietary preferences have low variability over time and did not change over seasons (Voigt and Kelm, 2006; Voigt *et al*., 2008; Voight, 2009; Streicker and Allgeier, 2016). This information was also supported by our direct observations, which showed that the common vampire bats can use memory and/or sensory cues to repeatedly feed on the same group of capybaras (Groger and Wiegrebe, 2006; Bahlman and Kelt, 2007, WebVideos S1 and S2).

Common vampire bats were not captured in studies in preserved areas of the mainland (Picinguaba, Mambucada e Santa Virgínia), which are near Anchieta Island (Supporting information S6, Fig. 1). This evidence corroborates the hypothesis that the percentage of capture of common vampire bats is usually low in natural forested habitats where potential prey are scattered, and high in areas with high concentration of prey (Turner, 1975; Bobrowiec *et al.*, 2015). Also, the number of individuals mist-nested varied on island nearby that have different history of human occupation (Supporting information S6, Fig. 1) indicating that common vampire bats may respond according to different type and intensity of anthropogenic disturbance (Streicker and Allgeier, 2016, Gonçalves *et al.*, 2017).

The current native mammal fauna on the Anchieta Island was quite impoverished due to its isolated location, as well as past human impact (Bovendorp and Galetti, 2007, Souza *et al.*, 2019, Supporting information S1). There are no previous studies about the occurrence and/or diet of common vampire bats on the island in the beginning of the last century (Garbino *et al.*, 2016, Muylaert *et al.*, 2017). However, previous information on human occupation on island (Guillaumon *et al.*, 1989) and record of common vampire bats in the mainland and island nearby Anchieta (Garbino *et al.*, 2016), led us to believe that domestic animals present there (especially cattle, pigs and dogs) were the main prey and responsible for maintaining a viable population of the species on island. When the island became a state park in 1977, all the domestic animals were removed (Guillaumon *et al.*, 1989) and forced the population of common vampire bats to leave or to reach very low density. After the species introductions in 1983, capybaras underwent explosive population growth due to food availability and absence of predators, and became an abundant and reliable source of blood for common vampire bats on Anchieta Island. The new scenario allowed common vampire bats to return to the island and/or increase the population densities. An alternative hypothesis is that common vampire bats never existed on the island and the new scenario created after species introductions allowed the species to colonize then.

The extent to which common vampire bats can shift to new food sources is poorly understood, but the degree to which they exhibit dietary shifts and how these feeding strategies respond to human activity, can be an indicator of community-level responses to environmental changes (Bolnick *et al.*, 2002; Layman *et al.*, 2007; Gonçalves *et al.*, 2017). The common vampire bat needs to feed every night (Freitas *et al.*, 2003), and prey that are dispersed or free to walk are more difficult to attack (Delpietro, 1989). Capybaras present a more predictable and constant food source on Anchieta Island, as they feed on grasses in open areas during the night, and are larger than coatis, reinforcing our conclusion that capybaras are the most attractive and reliable food source for common vampire bats in the study area.

The large biomass of capybaras on Anchieta Island and their predictable behavior, make them easy to find and more accessible than other potential prey for the common vampire bat. As the species was introduced to Anchieta Island only 36 years ago, this suggests a rapid prey shift due to anthropogenic disturbances, which has allowed common vampire bats to successfully exploit them. The shift from a livestock-based diet to introduced species poses interesting questions for common vampire bat health and behavior. Blood from translocate species might affect common vampire bats directly through differences in nutritional quality and exposure to new diseases, as detected in some bat individuals from Anchieta Island in which leptospirosis was serologically confirmed (Aires, 1998). Beyond that, three individuals of introduced capybaras were confirmed died from bat rabies viruses (RABV) in 2020 (PS Moreira, unpublished data) and we suggest, then, a periodic monitoring of bat rabies viruses (RABV) in common vampire bat populations on Anchieta Island and areas nearby, in order to quantify the magnitude of the outbreak area and develop strategies for controlling viruses, especially considering that the island and areas nearby is frequently visited by tourists.

In summary, stable isotope analysis is a useful tool for studying prey choice because it integrates information across wide time spans when quantified from tissues with slow turnover such as hair (Peterson and Fry, 1987), which in vampire bats, represent prey choice over 4–6 months prior to sampling (Voigt and Kelm, 2006). Our results indicate that, in the absence of livestock and domestic animals, vampire bats on Anchieta Island feed primarily on capybara, which is consistent with the bats having a preference for abundant species. The results are context-dependent and strongly influenced by: (1) the extirpation of domestic animals (2) the high abundance of this prey species, that is the highest mean biomass on the island; (3) the predictable foraging behavior of capybaras in open areas.

## Supplementary Material

**Supporting Information S1.** Introduced species and current population size and density of mammals [adapted from (Bovendorp and Galetti, 2007)] on the Anchieta island and their respective biomass [body mass according to Gonçalves et al. (2018)].

**Supporting Information S2.** Bat species recorded on the Anchieta Island, southeastern Brazil.

**Supporting Information S3.** δ^13^C and δ^15^N values of common vampire bats (*D. rotundus*) and potential prey (*H. hydrochaeris* and *N. nasua)* on Anchieta Island, southeastern Brazil, including the number of samples analyzed (N).

**Supporting Information S4.** Capture and sampling of common vampire bats (*D. rotundus*) and potential prey (1) capybaras (*H. hydrochaeris*) and (2) coati (*N. nasua)* on Anchieta Island, southeastern Brazil.

**Supporting Information S5.** Number of events of common vampire bats (*Desmodus rotundus*) feeding on capybaras (*Hydrochoerus hydrochaeris*) on Anchieta Island, southeastern Brazil.

